# Untamed: Unconstrained Tensor Decomposition and Graph Node Embedding for Cortical Parcellation

**DOI:** 10.1101/2024.01.05.574423

**Authors:** Yijun Liu, Jian Li, Jessica L. Wisnowski, Richard M. Leahy

**Affiliations:** Ming Hsieh Department of Electrical and Computer Engineering, University of Southern California, Los Angeles, CA, USA; Athinoula A. Martinos Center for Biomedical Imaging, Massachusetts General Hospital and Harvard Medical School, Charlestown, MA, USA; Center for Neurotechnology and Neurorecovery, Department of Neurology, Massachusetts General Hospital and Harvard Medical School, Boston, MA, USA; Radiology and Pediatrics, Division of Neonatology, Children’s Hospital Los Angeles, Los Angeles, CA, USA; Keck School of Medicine, University of Southern California, Los Angeles, CA, USA

**Keywords:** Cortical parcellation, Resting-state fMRI, Temporal synchronization, Tensor decomposition, Graph representation learning

## Abstract

Cortical parcellation is fundamental to neuroscience, enabling the division of cerebral cortex into distinct, non-overlapping regions to support interpretation and comparison of complex neuroimaging data. Although extensive literature has investigated cortical parcellation and its connection to functional brain networks, the optimal spatial features for deriving parcellations from resting-state fMRI (rsfMRI) remain unclear. Traditional methods such as Independent Component Analysis (ICA) have been widely used to identify large-scale functional networks, while other approaches define disjoint cortical parcellations. However, bridging these perspectives through effective feature extraction remains an open challenge. To address this, we introduce *Untamed*, a novel framework that integrates unconstrained tensor decomposition using NASCAR to identify functional networks, with state-of-the-art graph node embedding to generate cortical parcellations. Our method produces near-homogeneous, spatially coherent regions aligned with large-scale functional networks, while avoiding strong assumptions like statistical independence required in ICA. Across multiple datasets, Untamed consistently demonstrates improved or comparable performance in functional connectivity homogeneity and task contrast alignment compared to existing atlases. The pipeline is fully automated, allowing for rapid adaptation to new datasets and the generation of custom parcellations. The atlases derived from the Genomics Superstruct Project (GSP) dataset, along with the code for generating customizable parcel numbers, are publicly available at https://untamed-atlas.github.io.

## 1 Introduction

Elucidating the macrostructure of the human brain remains a cornerstone in neuroscience. From the early work of Brodmann though Talairach through the most recent efforts (Brodmann, 1909; Tzourio-Mazoyer et al., 2002; Yeo et al., 2011; Glasser et al., 2016; Schaefer et al., 2018; Eickhoff et al., 2018; Han et al., 2020; Joshi et al., 2022), a key aim has been identifying the parcels – the relatively homogenous mesoscale computational units that together give rise to the large-scale neural networks of the human brain.

The most common approach to elucidating the intrinsic macrostructure involves resting-state functional magnetic resonance imaging (rsfMRI). By recording blood-oxygen-level-dependent (BOLD) signals without task stimuli, rsfMRI proved instrumental in deciphering the human brain’s intrinsic functional brain networks (Allen et al., 2014; Beckmann et al., 2005; Biswal et al., 1995). From these, investigators have derived both group-level (Yeo et al., 2011; Craddock et al., 2012; Shen et al., 2013; Gordon et al., 2016; Schaefer et al., 2018; Yan et al., 2023) and individual (Chong et al., 2017; Kong et al., 2019) parcellations of the human brain. However, until recently, resting-state networks have been constrained by statistical independence or orthogonality, which are inherent to independent component analysis (ICA) or principle component analysis (PCA) approaches. Yet, these constraints may not be consistent with the intrinsic functional brain networks (Harris and Mrsic-Flogel, 2013; Rockland, 2019).

We have developed a new way for identifying overlapping and non-orthogonal functional networks using a Nadam-Accelerated SCAlabe and Robust (NASCAR) tensor decomposition method (Li et al., 2021, Li et al., 2023). The aim of this study is to determine whether cortical parcellations, derived from NASCAR networks, would be more robust when compared to prior parcellations. Briefly, NASCAR computes a three-way tensor decomposition of temporally aligned rsfMRI data across subjects (Joshi et al., 2018; Akrami et al., 2019) without imposing implausible constraints, such as statistical independence or orthogonality (Smith et al., 2014). It facilitates the identification of functional networks that may more authentically represent brain activity patterns by avoiding the need to impose spatial and/or temporal independence between networks. The networks identified by NASCAR have been shown to be, to a large degree, consistent with those found using ICA but with subtle differences that are consistent with other findings in the literature, such as subsystems of the default-mode network found using seed-based clustering and Bayesian methods (Andrews-Hanna et al., 2010; Andrews-Hanna et al., 2014; Buckner and DiNicola, 2019; Harrison et al., 2020). We also showed that the networks found using NASCAR could be used to better classify subjects with neurological disorders (Li et al., 2023).

Based on the brain networks identified by NASCAR, we construct a graph and apply a graph node embedding method, NetMF (Qiu et al., 2018), to generate cortical parcellations. This integration of tensor decomposition and graph node embedding enables the production of parcellations that are functionally homogeneous, computationally efficient, and adaptable to a wide range of datasets. Unlike parcellation methods that rely on manual input (e.g., (Glasser et al., 2016; Joshi et al., 2022)), our pipeline is fully automated, enabling researchers to seamlessly apply the framework to new datasets with a customizable number of parcels. This automation facilitates broader exploration of neuroscience questions, such as examining the influence of functional network spatial maps on the resulting parcels and determining the optimal number of parcels, as demonstrated in our experiments. Our approach, which we term *Untamed* (**Un**constrained **t**ensor decomposition-based **a**ctivation **m**aps and emb**ed**ding), delivers performance that is comparable to or exceeds widely used parcellations (e.g., (Schaefer et al., 2018)) across two critical metrics: resting-state functional connectivity homogeneity and task contrast alignment. By addressing methodological challenges in determining spatial features and linking cortical parcellation with functional networks, Untamed provides a robust framework for advancing research in neuroscientific studies. Both the parcellations derived from the Genomics Superstruct Project (Holmes et al., 2015) dataset and the code to generate customized atlases are available publicly at https://untamed-atlas.github.io.

## 2 Material and methods

### 2.1 Overview

We derived cortical parcellations using rsfMRI data from 1428 subjects from the Genomics Superstruct Project (GSP) dataset (Holmes et al., 2015). The overall pipeline is depicted in Fig. 1. The procedure included a temporal alignment using BrainSync (Akrami et al., 2019; Joshi et al., 2018), a 3-way tensor decomposition using NASCAR (Li et al., 2021, Li et al., 2023), graph construction from spatial maps followed by graph node embedding using NetMF (Qiu et al., 2018), and finally *k*-means clustering to obtain the parcellations. We evaluated the parcellation performance on the Yale rsfMRI (Lee et al., 2022), Human Connectome Project (HCP) (Van Essen et al., 2012; Glasser et al., 2013), and the multi-domain task battery (MDTB) (King et al., 2019; Zhi et al., 2022) datasets, with varied acquisition protocols and data modalities.

**Fig. 1.**
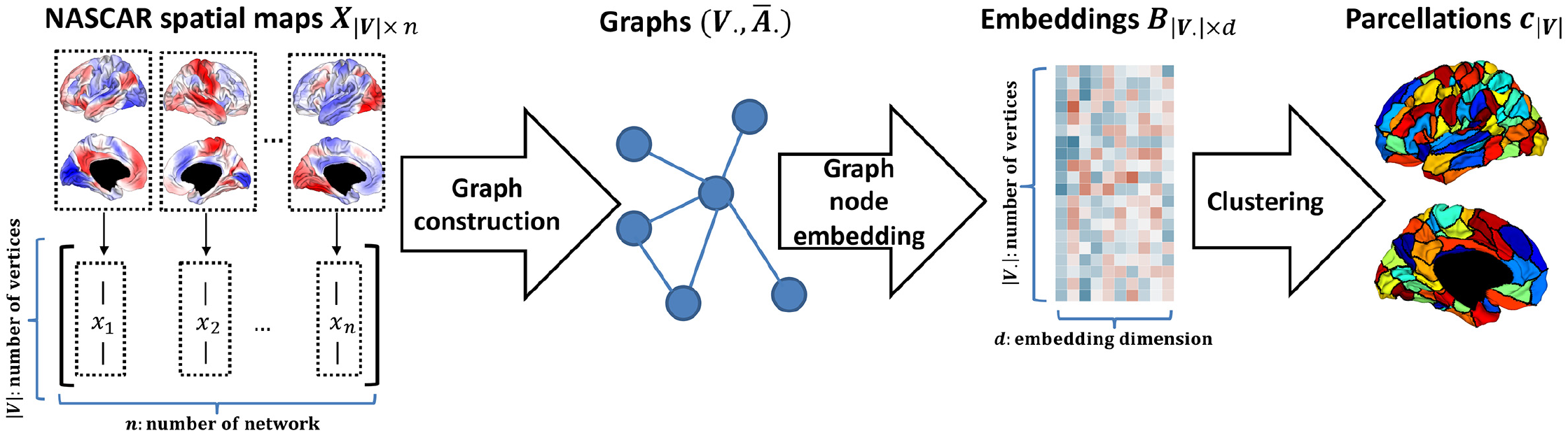
Pipeline to generate *Untamed* parcellations. See section 2.3-2.5 for the details of NASCAR spatial maps generation, graph construction, node embedding, and clustering.

### 2.2 Datasets

#### 2.2.1 The Genomics Superstruct Project (GSP) dataset

We utilized the Genomics Superstruct Project (GSP) (Holmes et al., 2015) dataset for atlas generation. The dataset contains 3T resting-state fMRI (rsfMRI) scans from 1570 subjects (665 males, 905 females, age between 22 and 35). Each subject underwent either one or two rsfMRI scans. Of the 1570 participants, 1139 completed two scans, while 431 completed only a single scan. The rsfMRI data were acquired with *TR* = 3s, and a 3 mm isotropic spatial resolution. Each session lasted 6 min and 12 s, yielding 124 time points. The structural data associated with each subject consisted of a single 1.2 mm isotropic scan.

Data preprocesssing was conducted using the publicly available pipeline from the CBIG repository (https://github.com/ThomasYeoLab/Standalone_CBIG_fMRI_Preproc2016), configured to closely align with methods described in (Li et al., 2019; Schaefer et al., 2018; Yan et al., 2023). Each subject’s T1-weighted image was registered to the standard MNI 2mm space. Key preprocessing steps of fMRI data included slice time correction, motion correction, censoring of outlier volumes, signal regression of white matter, ventricular signals, whole brain signal, and bandpass filtering (0.009 Hz – 0.08 Hz). Alignment between fMRI and structural images was performed using boundary-based registration (Greve and Fischl, 2009). The preprocessed fMRI were projected to the Freesurfer’s fsaverage6 space and smoothed using a Gaussian kernel with a 6 mm full-width half-maximum (FWHM). Subjects with at least one rsfMRI session that passed quality control checks in the preprocessing pipeline were included in subsequent experiments. This resulted in a final sample of 1428 subjects (603 males, 825 females), of which 1034 had two scans and 394 had one scan.

#### 2.2.2 The Human Connectome Project (HCP) dataset

We utilized the minimally preprocessed 3T rsfMRI data from 1000 subjects (466 males, 534 females, age between 22 and 35) from the Human Connec tome Project (HCP) (Glasser et al., 2013; Van Essen et al., 2012) for the assessment of parcellations. The rsfMRI data were acquired with *TR* = 0.72 s, *TE* = 33.1 ms, and a 2 mm isotropic resolution and co-registered onto a common atlas in MNI space (Glasser et al., 2013). We used the scans acquired in the LR phase encoding direction. Each session ran 15 minutes with 1200 time points. The data were resampled onto the cortical surface extracted from each subject’s T1-weighted MRI and co-registered to a common surface (Glasser et al., 2013). No additional spatial smoothing was applied other than the 2 mm FWHM isotropic Gaussian smoothing in the minimal preprocessing pipeline (Glasser et al., 2013).

In addition to rsfMRI data, we utilized subject-wise task activation z-score maps in fs_LR 32K surface space from the HCP dataset (Barch et al., 2013) for evaluation. These maps covered seven task domains: working memory, gambling, motor, language, social, relational, and emotion. Task contrast maps were derived from 3T task fMRI data acquired with the same parameters as the rsfMRI data, except for the duration which varied based on the particular task. We included all available contrasts and incorporated data from 1,006 subjects with complete task datasets. Furthermore, group-level task activation z-score maps for the same tasks were also employed for the analysis of optimal parcel numbers.

#### 2.2.3 Yale resting-state fMRI dataset

We used the Yale rsfMRI dataset (Lee et al., 2022) as an independent dataset for assessing RSFC homogeneity (Schaefer et al., 2018). The dataset comprises 27 subjects (11 males, 16 females, aged between 22 and 31). Each subject underwent two rsfMRI scans, each with *TR* = 1s, *TE* = 30ms, and a 2 mm isotropic resolution. The total duration of each session was 6 minutes 50 seconds, encompassing 410 frames. We preprocessed the data using fMRIPrep (Esteban et al., 2019), which sampled the fMRI data onto the fs_LR 32K surfaces compatible with the HCP dataset. Four subjects’ data were not successfully preprocessed by the fMRIPrep pipeline due to corrupted files and were consequently excluded from the evaluation. Following the recommendations from the curators of the dataset, we discarded the first 10 seconds of each scan and temporally concatenated the rsfMRI data of the two sessions for each subject.

#### 2.2.4 Multi-domain task battery (MDTB) dataset

We employed the multi-domain task battery (MDTB) dataset (King et al., 2019; Zhi et al., 2022) for evaluating the alignment between parcels in the atlases and areas of activation in the task activation maps. The MDTB dataset contains task fMRI for 24 healthy subjects (8 males, 16 females, mean age 23.8 years old) conducting 26 tasks, including motor, language, and social domains. The fMRI data were acquired on a 3T Siemens Prisma scanner with *TR* = 1s, 3 mm slice thickness, and 2.*L* × 2.*L* mm^2^ in-plane resolution. The contrast maps were derived using general linear modeling (GLM) based on the task designs and are available from the database, already resampled to the fs_LR 32K space.

### 2.3 Tensor-based identification of Brain Networks using NASCAR and BrainSync

For each subject, if two sessions of rsfMRI data were available, the time series from the two sessions were concatenated temporally. If only one session was available, the session was duplicated and concatenated. The resulting multisubject rsfMRI data were then organized as a three-way tensor (space × time × subject). In order to identify a low-rank model via tensor decomposition, the data needed to be both temporally and spatially aligned. As described in the preprocessing pipeline for the GSP dataset, inter-subject spatial alignment was achieved using standard image registration methods, mapping each subject’s data to the standard fsaverage6 surface space. However, while spatial alignment ensured that brain regions corresponded across subjects, the rsfMRI time series remained independent for each subject. To address this we assume that subjects share a similar connectivity structure as reflected in the pairwise correlation patterns between brain regions.We can then apply an orthogonal transformation to temporally align or synchronize the data across subjects. This temporal alignment is achieved using the BrainSync transform (Joshi et al., 2018). BrainSync computes an orthogonal transformation between fMRI recordings from a pair of subjects such that the sum of squared errors between their aligned time series is minimized. As a result, the time series in homologous brain locations will be highly correlated after the BrainSync transform is applied (and perfectly so if the two datasets have identical correlations). A multi-subject extension, as described in (Akrami et al., 2019), minimizes the squared error from all subjects to an automatically generated group template. This method was used in this study to align all subjects for generating the cortical parcellation.

After temporal synchronization, we formed a third-order tensor ***X*** ∈ ℝ^|***V***|×*T*×*S*^ from the spatially aligned and temporally synchronized rsfMRI data, where ***V*** is the set of cortical vertices with a cardinality |***V***| ≈ 75*K, T* = 240 is the number of time points, *S* = 1428 is the number of subjects used to generate the atlas. We then applied NASCAR to decompose the group rsfMRI data into a set of brain networks common to all subjects using a Canonical Polyadic model (Kolda and Bader, 2009; Cichocki et al., 2015; Li et al., 2021, Li et al., 2023)

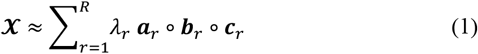

where each outer product *λ*_*i*_ ***a***_*i*_ ∘ ***b***_*i*_ ∘ ***c***_*i*_ represents a brain network: ***a***_*i*_ ∈ ℝ^|***V***|^ are the spatial network maps, ***b***_*i*_ ∈ ℝ^*T*^ their (synchronized) temporal dynamics, and ***c***_*i*_ ∈ ℝ^*S*^ the subject participation level for the *i*^th^ network; *λ*_*i*_ represents the network magnitude, indicating the relative activity level with respect to other networks. In contrast to the commonly used blind source separation techniques such as PCA or ICA, NASCAR imposes neither orthogonality (as with PCA) (Smith et al., 2014) nor statistical independence (as with ICA) (Beckmann et al., 2005). The shared spatial network maps {***a***_*i*_} can be overlapped and correlated (Li et al., 2023). Based on our previous work that examined stability and reproducibility across data sets, we chose a rank *R*=50 (Li et al., 2023). We performed NASCAR decomposition on the whole GSP dataset and obtained a set of 50 spatial network maps, which formed a spatial feature matrix ***X*** ∈ ℝ^|***V***|×50^ (|***V***| ≈ 75*K*). This matrix provided a 50-dimensional feature vector for each vertex. Specifically, ***X***_*ij*_ represented the participation level at the *i*^th^ vertex in the *j*^th^ network map.

### 2.4 Graph construction from NASCAR spatial maps

We computed the Pearson correlation matrix ***A*** = corr(***X***) ∈ ℝ^|***V***|×|***V***|^ from the feature matrix ***X*** as a measure of similarity between feature vectors of each pair of vertices. We then computed an adjacency matrix, following (Ng et al., 2001), using the Gaussian kernel 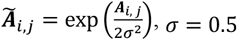. Our goal was to produce contiguous parcels, and as such, we were not interested in long-range correlations. Therefore, we generated a graph where connections were restricted using an *nb*-hop spatial neighborhood constraint defined on the fs_LR 32K surface mesh. Specifically, we defined 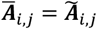 for all (*i, j*) for which *i* is within *nb* hops of *j* and zero otherwise. In a related approach, (Craddock et al., 2012) used a *j*-hop neighborhood, retaining nearest neighbors only.

However, we found that a larger *nb* could produce higher RSFC homogeneity (see Evaluation section). Consequently, rather than fixing this parameter, we treated it as a hyperparameter within the algorithm, optimized using the GSP dataset. Details of the hyperparameter selection process are provided in Section 2.5.

Using the above approach, we obtained one sparsely connected (*m*-hop) graph for each hemisphere: 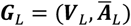 and 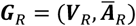, where ***V***_*L*_, ***V***_*R*_ were the set of cortical vertices and 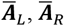 were the *nb*-hop filtered adjacency matrices of the graphs ***G***_*L*_, ***G***_*R*_, for the left and the right hemispheres, respectively.

### 2.5 Graph node embedding and clustering

We adopted the method outlined in NetMF (Qiu et al., 2018) to embed vertices in a lower dimensional space with more representative features that can be used for clustering. This approach (Qiu et al., 2018) established an equivalence between the widely-used DeepWalk algorithm (Perozzi et al., 2014) and matrix factorization techniques. Briefly, DeepWalk traverses vertices in the graph using a random walk. At each vertex during traversal, DeepWalk considers neighboring vertices as co-occurrent “positive” pairs as well as random node pairs outside the neighborhood as “negative samples.” It then applies a skip-gram model to train on the collected samples to derive the final embeddings. NetMF approximates DeepWalk by factorizing the following matrix (Qiu et al., 2018):

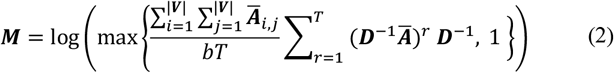

where ***D*** is the degree matrix of the graph; log(·) and max{·, *j*} are element-wise logarithmic and maximum operations, respectively; *b* denotes the number of negative samples; and *T* is the window length over which nodes considered as co-occurrent positive pairs. We found that in practice the method was not sensitive to the choice of these two hyperparameters, which had little impact on the averaged training subjects’ RSFC homogeneity (see evaluation metrics section). We set b = *j*, the default value in the original NetMF paper. We computed the ***M*** matrices for left hemisphere from 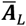 and for right hemisphere from 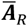 separately. We used singular value decomposition (SVD) to obtain the final embeddings (Qiu et al., 2018). Specifically, let 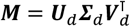 where **Σ**_*d*_ and ***U***_*d*_ being the *d* largest singular values and their associated left singular vectors. The final embeddings were computed as 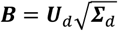 for the left (***B***_*L*_) and right hemisphere (***B***_*R*_) separately, with *d* being the embedding dimension.

To obtain the final cortical parcellation, we applied *k*-means clustering to the embedding vectors formed by the columns of ***B***_*L*_ and ***B***_(_ and varied the number of classes *k* to match the number of desired parcels in the parcellation. For each value of *k*, we ran *k*-means clustering with 500 different random initializations with a maximum of 20000 iterations and selected the result that yielded the smallest cost across trials as the final output.

The graph construction and embedding procedure involved fine tuning the following parameters: spatial neighborhood constraint *nb*, the window length *T*, and the embedding dimension *d*. Specifically, *nb* was varied across{1, 5, 10, 15, 20, .…, 60}, *T* across {1, 7, 11, 15}, and *d* across {128, 256}. The finetuning process aimed to maximize the weighted average RSFC homogeneity. Hyperparameter combinations were selected based on the mean weighted average RSFC homogeneity across GSP subjects, which served as a summary statistic to assess performance on the training set.

The following parameter combinations were selected based on the fine tuning results:

- Embedding dimension = 128
- Neighborhood size (*nb*) and window length (*T*):
  ∘ *nb* = 55, *T* = 7 for parcel number ∈ (0, 200]
  ∘ *nb* = 40, *T* = 15 for parcel number ∈ (200, 300]
  ∘ *nb* = 35, *T* = 15 for parcel number ∈ (301, 400]

### 2.6 Evaluation metrics

#### 2.6.1 Resting-state functional connectivity (RSFC) homogeneity

Each subject’s RSFC was computed as the Pearson correlation of rsfMRI time series between all pairs of cortical vertices. A Fisher z-transform was subsequently performed to obtain z scores from the correlation values. Similar to the evaluation procedure in (Joshi et al., 2022; Schaefer et al., 2018), a parcel-wise homogeneity score *ρ*_*i*_ was computed as the averaged RSFC values within the *i*^th^ parcel. To obtain a global homogeneity measure for a single subject, a weighted average *ρ* of each parcel’s homogeneity scores was then computed by accounting for different cluster sizes:

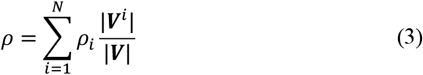

where |***V***^*i*^| is the number of vertices in parcel *i*, and *N* is the total number of parcels; |***V***| is the total number of cortical vertices. For conciseness, we refer to this RSFC weighted average homogeneity as “homogeneity” hereafter. The homogeneity was computed for each test subject separately. To quantitatively compare the homogeneity of Untamed to that of other atlases, we performed a paired-sample t-test. Effect size was reported using Cohen’s *d* measure.

#### 2.6.2 Alignment with task contrasts

The degree to which the regions delineated in a particular parcellation reflect functional specialization was assessed by computing the variance 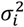 of the task contrast within each parcel. The better the parcellation delineates regions of functional homogeneity, the lower the variance of task contrasts within each parcel. Similar to the procedure in (Schaefer et al., 2018; Joshi et al., 2022; Yan et al., 2023), the variance metric was first computed for each parcel, and then a weighted average was computed, accounting for parcel size differences, as:

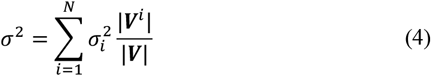

We refer to this weighted average task contrast variance as “task variance” hereafter. The variance was computed separately for each task contrast and for each subject. These variances were first averaged across contrasts within each task and then averaged across tasks, consistent with the approach in (Yan et al., 2023). Statistical significance was assessed using paired-sample t-test to compare the task variance between different parcellations.

### 2.7 Comparison with existing parcellations

We compared our Untamed parcellations to an extensive set of 14 atlases listed in Table 1. Because comparison of the evaluation metrics defined above was only meaningful when comparing parcellations of the same parcellation resolution, in each case, we matched the number of parcels found using Untamed to the number in the left and right hemispheres for each baseline comparison.

**Table 1.** List of atlases included in the comparison. Type denotes the modality of data the atlas is constructed from. “Functional”: the atlas is only based on fMRI; “Hybrid”: the atlas is constructed from both functional MRI and anatomical information.

| Name | Type | # Clusters | Reference | Atlas Source |
| --- | --- | --- | --- | --- |
| Yeo-51<br>Yeo-114 | Functional | 51 (L: 26, R: 25)<br>114 (L: 57, R: 57) | (Yeo et al., 2011) | https://github.com/ThomasYeoLab/CBIG/tree/master/stable_projects/brain_parcellation |
| Power | Functional | 130 (L: 65, R: 65) | (Arslan et al., 2018) | https://biomedica.doc.ic.ac.uk/brain-parcellation-survey/ |
| USCBrain | Hybrid | 130 (L: 65, R: 65) | (Joshi et al., 2022) | https://github.com/ajoshiusc/bfp/blob/main/supp_data/USCBrain_grayordinate_labels_clean.mat |
| Shen | Functional | 200 (L: 102, R: 98) | (Shen et al., 2013) | https://biomedica.doc.ic.ac.uk/brain-parcellation-survey/ |
| Schaefer-100/200/300/400 | Functional | 100 (L: 50, R: 50)<br>200 (L: 100, R: 100)<br>300 (L: 150, R: 150)<br>400 (L: 200, R: 200) | (Schaefer et al., 2018) | https://github.com/ThomasYeoLab/CBIG/tree/master/stable_projects/brain_parcellation |
| Yan-100/200/300/400 | Functional | 100 (L: 50, R: 50)<br>200 (L: 100, R: 100)<br>300 (L: 150, R: 150)<br>400 (L: 200, R: 200) | (Yan et al., 2023) | https://github.com/ThomasYeoLab/CBIG/tree/master/stable_projects/brain_parcellation |
| Glasser | Hybrid | 360 (L: 180, R: 180) | (Glasser et al., 2016) | https://balsa.wustl.edu/study/show/RVVG |

The evaluations were conducted in the HCP fs_LR 32K surface space. Untamed, originally constructed in the fsaverage 6 surface space, was projected onto the HCP fs_LR 32K space. For other atlases that were originally constructed in spaces different from HCP fs_LR 32K (e.g., MNI152, fsaverage), we utilized versions of the atlases that had been resampled onto the HCP fs_LR 32K surfaces. These resampled versions were provided either by the original authors or by third parties, as detailed in Table 1. For the original two atlases proposed by (Yeo et al., 2011) that are spatially distributed across hemispheres with a relatively small number (7 and 17) of networks, we used the version provided in their GitHub repository (Table 1), where the distributed spatial networks were decomposed into local contiguous parcels (51 parcels for the 7 networks and 114 parcels for the 17 networks).

### 2.8 Ablation study of input spatial maps: NASCAR versus ICA

In the Untamed framework, the graph was constructed using correlations between features representing the participation of each vertex in the NASCAR spatial network maps. However, the input spatial maps are not limited to NASCAR and can be derived from any network decomposition method. To investigate whether the non-orthogonal spatial maps generated by NASCAR lead to better performance compared to spatial maps from other methods, all steps in the pipeline were kept consistent, with the only variation being the input spatial maps. For this study, ICA spatial maps provided by HCP were used as an alternative to NASCAR maps. Both the ICA and NASCAR maps consisted of 50 networks, with no additional selection or preprocessing applied. The pipeline’s hyperparameters—including the number of spatial neighborhood constraints, the window length in the NetMF algorithm, and the embedding dimension—were optimized specifically for the ICA maps and explored within the same range as those used for NASCAR. RSFC homogeneity was evaluated on the HCP rsfMRI dataset consisting of 1000 test subjects using the same procedure described in 2.6.1. To statistically compare the homogeneity values derived from the NASCAR and ICA spatial maps, a paired-sample t-test was conducted. This analysis was designed to evaluate how the choice of input spatial maps influences the performance of the Untamed framework.

We note that the NASCAR spatial maps that Untamed atlases generated from were derived from the independent GSP dataset in the fsaverage 6 surface space and subsequently projected onto the HCP fs_LR 32K surface space. In contrast, the ICA maps from HCP inherently benefit from dataset-specific fine-tuning and remain within the same surface space, avoiding potential information loss associated with the projection process.

### 2.9 Ablation study of graph embedding methods

We also compared Untamed with the spectral clustering described in (Ng et al., 2001), which applies a *k*-means clustering to the most significant eigenvectors of the normalized graph Laplacian. All steps in the learning procedure were identical except for the graph embedding, where we used NetMF. We applied both methods to the same graph constructed from NASCAR spatial maps as described in section 2.4. Parcellations generated from NetMF and graph Laplacian were subject to the same neighborhood constraints, embedding dimension, and k-means clustering hyperparameters with a range of cluster numbers. We evaluated weighted average homogeneity on the 1000 HCP rsfMRI dataset test subjects. A paired-sample t-test test was used to compare the homogeneity values obtained from the Laplacian eigenvectors used in spectral clustering and the NetMF method that was utilized in Untamed.

### 2.10 Ablation study of graph construction methods

In Untamed, we constructed the graph using the correlation between features representing the participation of each vertex in each of the NASCAR spatial network maps. Here, we conducted a comparison to the graph constructed from the widely used correlation map of RSFC, i.e. the correlation of the correlation between rsfMRI time series. Other steps were identical except for the construction of the graph. Parcellations generated from NASCAR spatial maps and RSFC correlation maps were subject to the same neighborhood constraints and embedding dimension. We evaluated weighted average homogeneity on the averaged RSFC from the 500 training subjects in the HCP dataset. A paired-sample t-test was employed to statistically compare homogeneity values obtained from NASCAR spatial maps and the correlation of RSFC.

### 2.11 Is there a (natural) optimal number of parcels that can be identified from rsfMRI data?

To address this question, we evaluated the performance of the Untamed atlas in identifying regions of relatively homogeneous functional activity, as measured by RSFC homogeneity (described in Section 2.6.1) and task contrast variance (described in Section 2.6.2), relative to random parcellations with an equal number of parcels on the HCP dataset. Random parcellations were employed as a null model to assess whether there was an optimal number of parcels for functional parcellation.

Specifically, we calculated the ratio of the metric scores obtained with the Untamed atlas to those from random parcellations for both RSFC homogeneity and task variance. Random parcellations were generated using a region-growing process implemented in the MNE-Python package (Gramfort, 2013). This process was repeated across a range of parcel numbers, from 1 to 500 parcels per hemisphere.

For each parcel number, 50 random parcellation realizations were generated with different initialization seeds. The mean weighted average metric scores across these 50 random parcellations were computed and used to calculate the performance ratio with respect to the Untamed atlas. These ratios were then plotted as a function of the number of parcels to investigate the relationship between parcel number and the performance of the Untamed atlas relative to the null model.

## 3 Results

### 3.1 Resting-state functional connectivity (RSFC) homogeneity

Fig. 2 (a) (Yale dataset) and Fig. 2(b) (HCP dataset) present the weighted average RSFC homogeneity as bar charts. Corresponding effect sizes, quantified by Cohen’s d, are reported in Table S1. In Fig. 2(a), results from the Yale dataset demonstrates that the Untamed method outperformed baseline methods across all tested parcel numbers with statistical significance (*p* < .001, *uncorrected*). Similarly, Fig. 2 (b) shows that on the HCP dataset, the Untamed method consistently achieved higher weighted average homogeneity compared to baseline methods, with most differences being statistically significant (*p* < .001), except for the 300-parcel case, where no statistically significant differences were observed between Untamed-300 and Schaefer-300 or Yan-300.

**Fig. 2.**
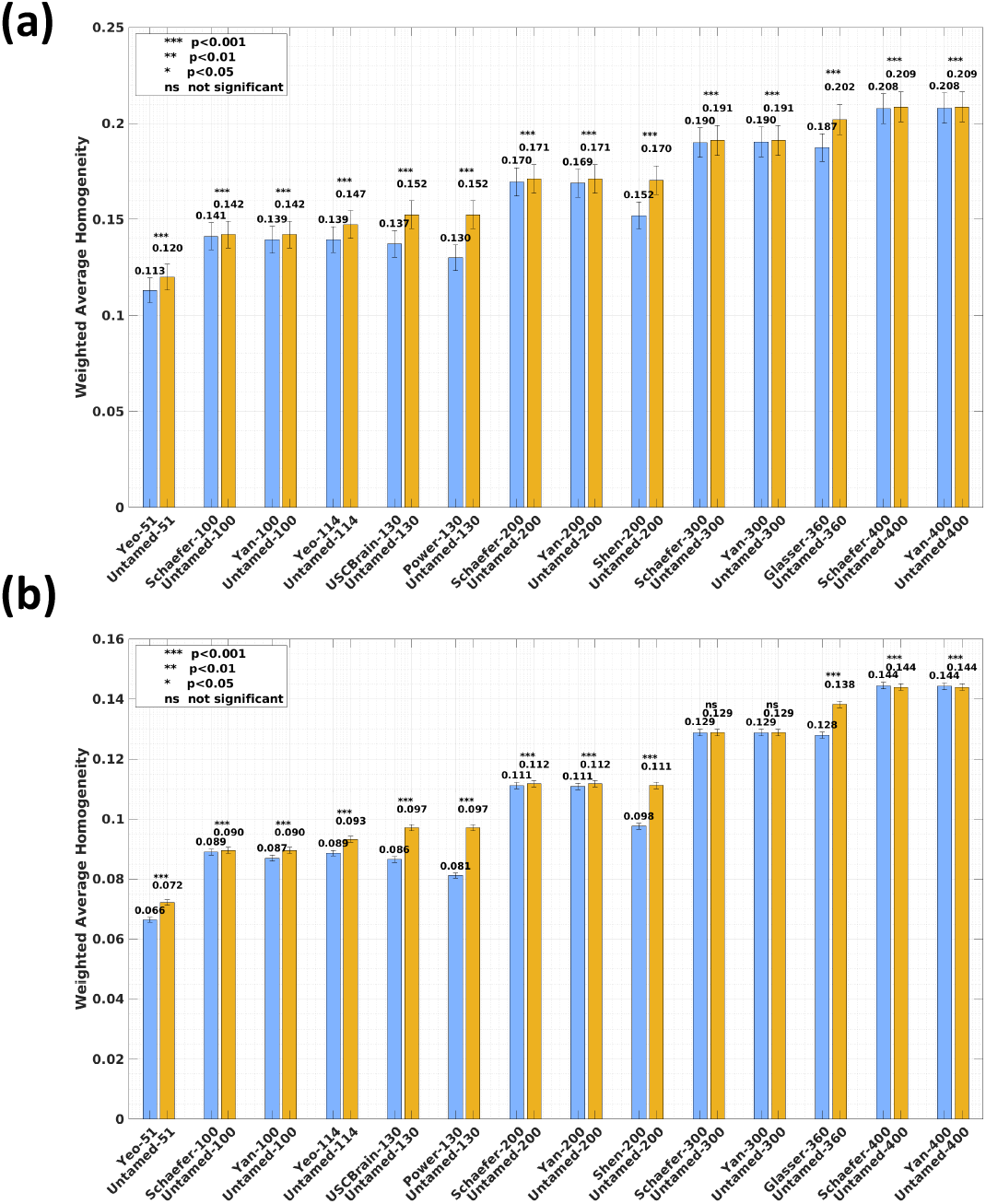
Weighted average RSFC homogeneity on the (a) Yale (b) HCP dataset. Each bar plot depicts the subject-wise RSFC weighted average homogeneity, averaged across test subjects, for each baseline and the Untamed atlas, with matched parcel numbers for the left and right hemispheres. The error bars represent the standard error across all subjects. Effect sizes are shown in Table S1.

An exception is noted in the 400-parcel case on the HCP dataset, where Schaefer-400 and Yan-400 achieve slightly higher homogeneity values than the Untamed method. However, the differences are negligible, with no variation up to the third decimal place and effect sizes of 0.0151 and 0.0085, respectively. Overall, these findings highlight the superior performance of the Untamed method, which generally produces more functionally homogeneous brain parcellations than other widely used approaches. This superiority was consistently observed across different datasets, parcel numbers, and parcellation schemes, emphasizing the robustness and effectiveness of the Untamed atlases.

Interestingly, despite the large sample size of the HCP dataset, no statistical significance is detected between Untamed-300 and either Schaefer-300 or Yan-300. This suggests negligible differences between these methods in the 300-parcel case. This observation is further explored in the Discussion section.

### 3.2 Alignment with task contrasts

The violin plots in Figure 3 (a, b) depict the distribution of task contrast variance (as defined in Eq. 4) for the Untamed method and baseline atlases with matching parcel numbers. Fig. 3 (a) represents results from the MDTB task dataset, while Fig. 3 (b) illustrates results from the HCP task dataset. Lower task contrast variances correspond to a smaller overall contrast variance per parcel, indicating better alignment of parcellation with functional task responses.

**Fig. 3.**
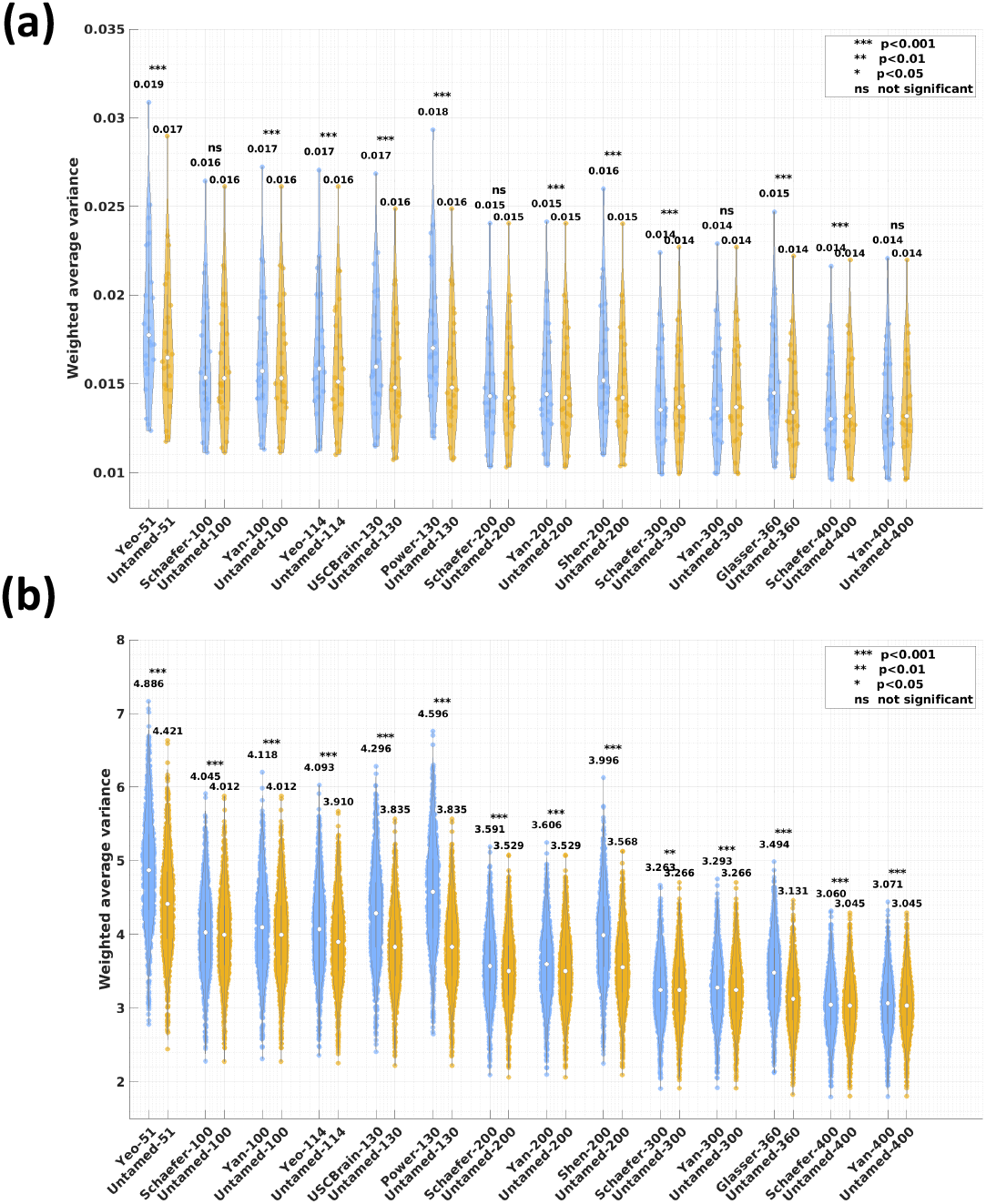
Weighted average task contrast variance evaluated on (a) MDTB (b) HCP test subjects. Each violin plot depicts the alignment with task contrast maps, computed for the baseline and the Untamed atlas, with matched parcel numbers for the left and right hemispheres.

The Untamed method demonstrates statistically significant improvements over baseline methods across almost all comparisons on the HCP dataset. The only exception is Schaefer-300, which shows a lower task contrast variance than its Untamed counterpart; however, this difference does not reach statistical significance after applying Bonferroni correction for multiple comparisons (raw *p* > .005). On the MDTB dataset, only Schaefer-300 and Schaefer-400 show better alignment than their Untamed counterparts with statistical significance. However, the differences are indistinguishable up to the third decimal place. For the remaining comparisons, Untamed either outperforms baseline methods with statistical significance or shows comparable performance (e.g., Schaefer-100/200, Yan-300/400, where no significant differences are observed).

Although Glasser-360 explicitly used HCP task fMRI data (in combination with structural data) in constructing the atlas, the variances computed on both the HCP and MDTB task data were significantly higher than those of Untamed.

Fig. 4 (a, b) illustrates the overlay of Schaefer-100 and Untamed-100 parcels on the HCP group average “story” contrast map from the language task. Regions near Broca’s area and other auditory regions demonstrate superior alignment of the Untamed parcels with contrast compared to Schaefer’s parcellation. Conversely, a counterexample is presented in Figures 6 (c, d), where the “relational_match” contrast reveals better alignment of task-active regions with the Schaefer parcellation. The weighted average variance of language_story contrast is 4.3892 for Untamed-100 and 5.4311 for Schaefer-100. Meanwhile, the weighted average variance of relational_match contrast is 8.4673 for Untamed-100 and 8.0539 for Schaefer-100 (lower variance indicates better alignment).

**Fig. 4.**
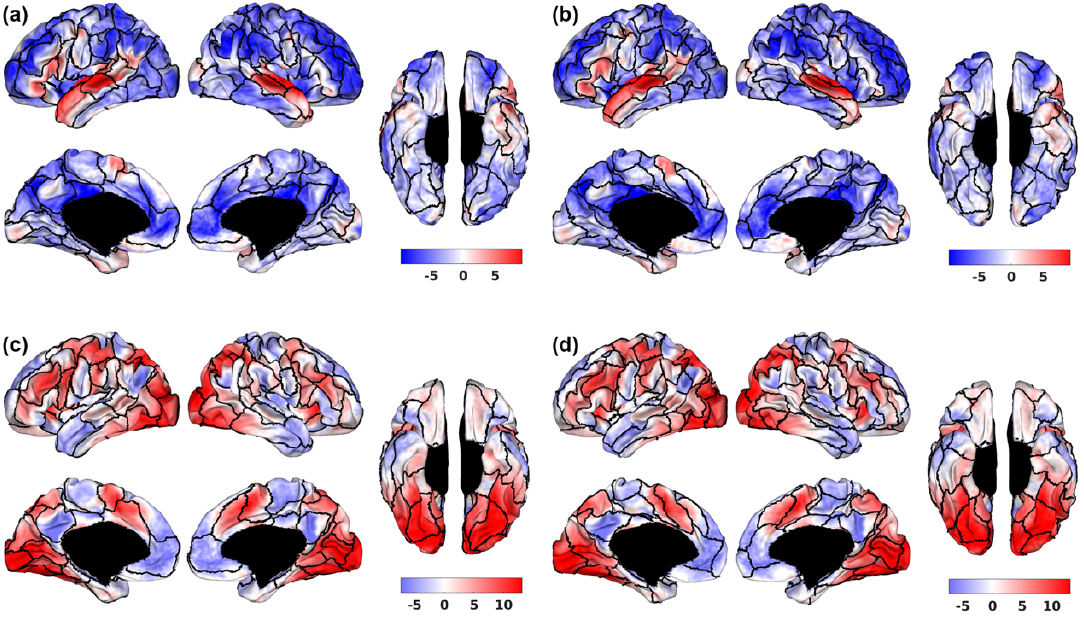
Example HCP group average task activation z-score maps for Schaefer-100 and Untamed-100. First row: language_story contrast overlaid on boundaries (black) of (a) Schaefer-100 (b) Untamed-100. Second row: relational_match contrast overlaid on (c) Schaefer-100 (d) Untamed-100.

Overall, the Untamed method consistently achieves lower weighted average variance compared to baseline methods across most parcel numbers and parcellation schemes. This result highlights the ability of the Untamed method to produce functionally homogeneous brain regions with reduced variability in functional task responses.

### 3.3 Ablation study of input spatial maps: NASCAR versus ICA

Fig. 5 shows the weighted average RSFC homogeneity of the 1000 subjects in the HCP dataset. The results indicates that despite the fact that using the ICA maps from HCP has the advantage of same-dataset hyperparameter fune-tining and remaining in the same surface space, using input spatial maps from NASCAR still consistently outperformed ICA across all four parcel numbers with statistical significance (*p* <.001).

**Fig. 5.**
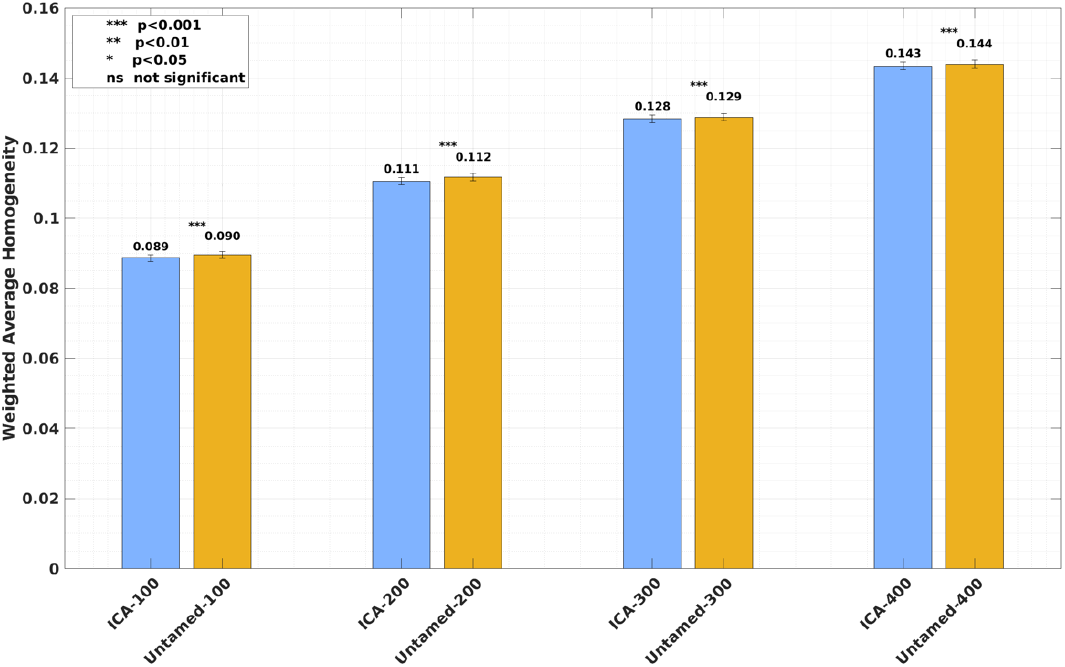
Weighted average RSFC homogeneity on the HCP dataset. Each bar plot depicts the subject-wise RSFC weighted average homogeneity, averaged across est subjects, for each atlas generated from ICA maps and the Untamed atlas (generated from NASCAR maps). Parcel numbers matched for the left and right hemispheres. The error bars represent the standard error across all subjects. The ICA baselines, derived from and optimized through hyperparameter tuning on the same HCP dataset, benefit from this dataset-specific tuning. In contrast, the Untamed atlas was generated using the independent GSP dataset with hyperparameters tuned on GSP. The ICA results represent the maximum homogeneity achievable under optimal hyperparameter settings for the HCP dataset.

**Fig. 6.**
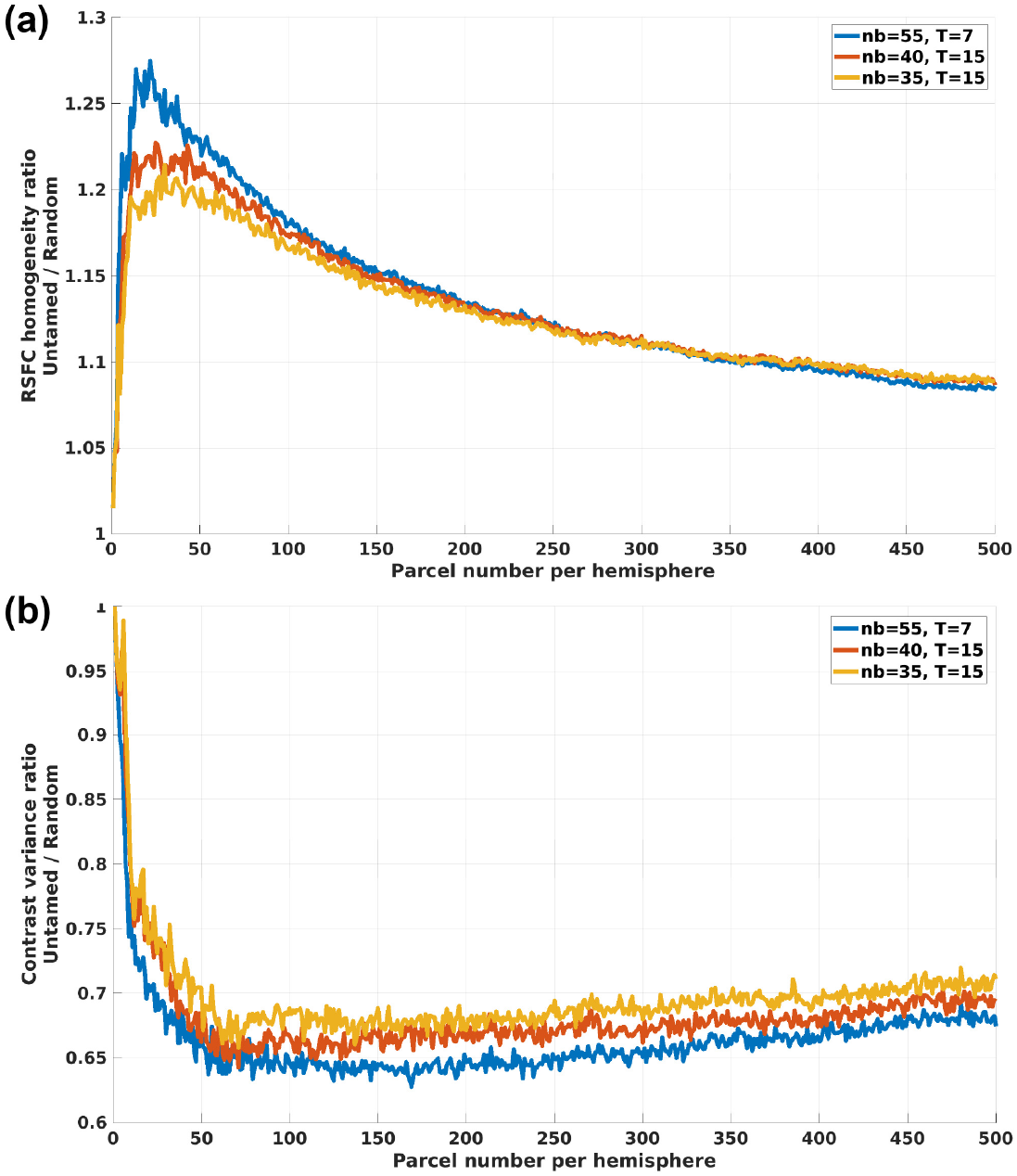
Ratios of three evaluation metrics comparing Untamed and random parcellations across 1 to 500 parcels per hemisphere: (a) Ratio of weighted average RSFC homogeneity: a comparison between Untamed parcellations and the mean values from 50 random parcellation trials (b) Ratio of weighted average task contrast variance: a comparison between Untamed parcellations and the mean values from 50 random parcellation trials.

### 3.4 Ablation study of graph embedding methods

In Fig. S1, we show parcellations obtained using graph node embedding (NetMF) prior to clustering as described above with results obtained using spectral clustering directly from the eigenvectors of the graph Laplacian (GLC). Again, we show the bar plots of Untamed and GLC-based atlases in RSFC weighted average homogeneity per HCP test. Among the 4 different numbers of parcels tested, NetMF-based Untamed outperforms the GLC-based one in 3 cases with statistical significance (*p* < .001). These results support the use of NetMF embedding in place of the more standard GLC approach.

### 3.5 Ablation study of graph construction methods

We also explored the effect of graph construction as described in Section 2.10 by comparing results using the NASCAR-based adjacency matrix with that computed using the correlation of the RSFC matrix. All other aspects of processing were identical. Fig. S2 shows the bar plots of RSFC weighted average homogeneity of NASCAR-based and RSFC-based methods. Evidently, the parcellations generated from the NASCAR-based adjacency matrix substantially outperformed those generated from Pearson-based adjacency in all cases (*p* < .001). This demonstrated an advantage of using the results of NASCAR tensor decomposition to identify spatial networks over directly using the correlation of RSFC.

### 3.6 Is there an optimal number of parcels?

Fig. 6 illustrates the ratio of RSFC homogeneity and task contrast variance between the Untamed atlas and random parcellations for the HCP task dataset. For RSFC homogeneity, a larger ratio indicates comparatively higher homogeneity for the Untamed atlas relative to its random parcellation counterpart. Conversely, for task contrast variance, a smaller ratio indicates better performance by the Untamed atlas.

Both curves exhibit a similar trend: the advantage of the Untamed atlas over random parcellations increases initially, peaks, and then diminishes as the number of parcels continues to grow. The optimal number of parcels varies between the two modalities. Fig. 6 (a) indicates that the greatest relative advantage of the Untamed atlas over random parcellations occurs at fewer than 50 parcels per hemisphere for the homogeneity metrics as evidenced by the peak. The contrast variance ratio in Fig. 6 (b) remains relatively flat and close to its minimum from approximately 50-150 parcels with the advantage relative to a random parcelation diminishing approximately montonically above 200 parcels.

## 4 Discussion

This paper introduced *Untamed*, a novel cortical parcellation scheme developed from population resting-state fMRI data. This scheme constructed spatially disjoint parcels by leveraging the overlapping and correlated brain networks identified by NASCAR tensor decomposition (Li et al., 2023, 2021). We compared Untamed to an extensive list of popular atlases and parcellation methods, as listed in Table 1. As noted in Section 3.1, the weighted average RSFC homogeneity among the HCP subjects revealed no statistically significant differences between Untamed-300 and Schaefer-300, nor between Untamed-300 and Yan-300. The large sample size of 1000 test subjects indicated that the differences in homogeneity between Untamed and these two atlases at this particular parcel resolution were minimal. Nonetheless, a more detailed examination of the per-parcel RSFC homogeneity revealed nuanced differences. We conducted an in-depth comparison between Untamed-300 and Schaefer-300. As illustrated in Fig. 7 (a) and (b), the overall spatial distribution of per-parcel RSFC homogeneity exhibited a broadly similar pattern across the cortex for both parcellations. Specifically, in both cases, parcels in the parietal and occipital lobes exhibit higher RSFC homogeneity compared to those in the frontal lobe. This may be attributed to the more functionally specialized regions in the parietal and occipital areas, which contribute to stronger intra-parcel connectivity, whereas the frontal lobe encompasses more heterogeneous and integrative functions. Despite these general similarities, visible differences in per-parcel RSFC homogeneity were observed across several cortical areas. Fig. 7 (c) quantifies these differences, highlighting variations in RSFC homogeneity values between parcels for the two parcellations. While the weighted average RSFC homogeneity reported in section 3.1 provides a global perspective on performance across the entire cortex, per-parcel RSFC homogeneity scores offer a finer-grained view. Notably, the per-parcel RSFC homogeneity scores between Untamed and Schaefer remained significantly more consistent compared to those between Untamed and a random parcellation (as described in Section 2.10). For easier visual comparison, we provide per-parcel homogeneity plots similar in style to Fig. 7, illustrating Untamed-100 vs. Schaefer-100 and Untamed-100 vs. Random-100 in Fig. S3 and S4. The random parcellation showed substantially less alignment, both in terms of parcel overlap and parcel-wise RSFC homogeneity.

**Fig. 7.**
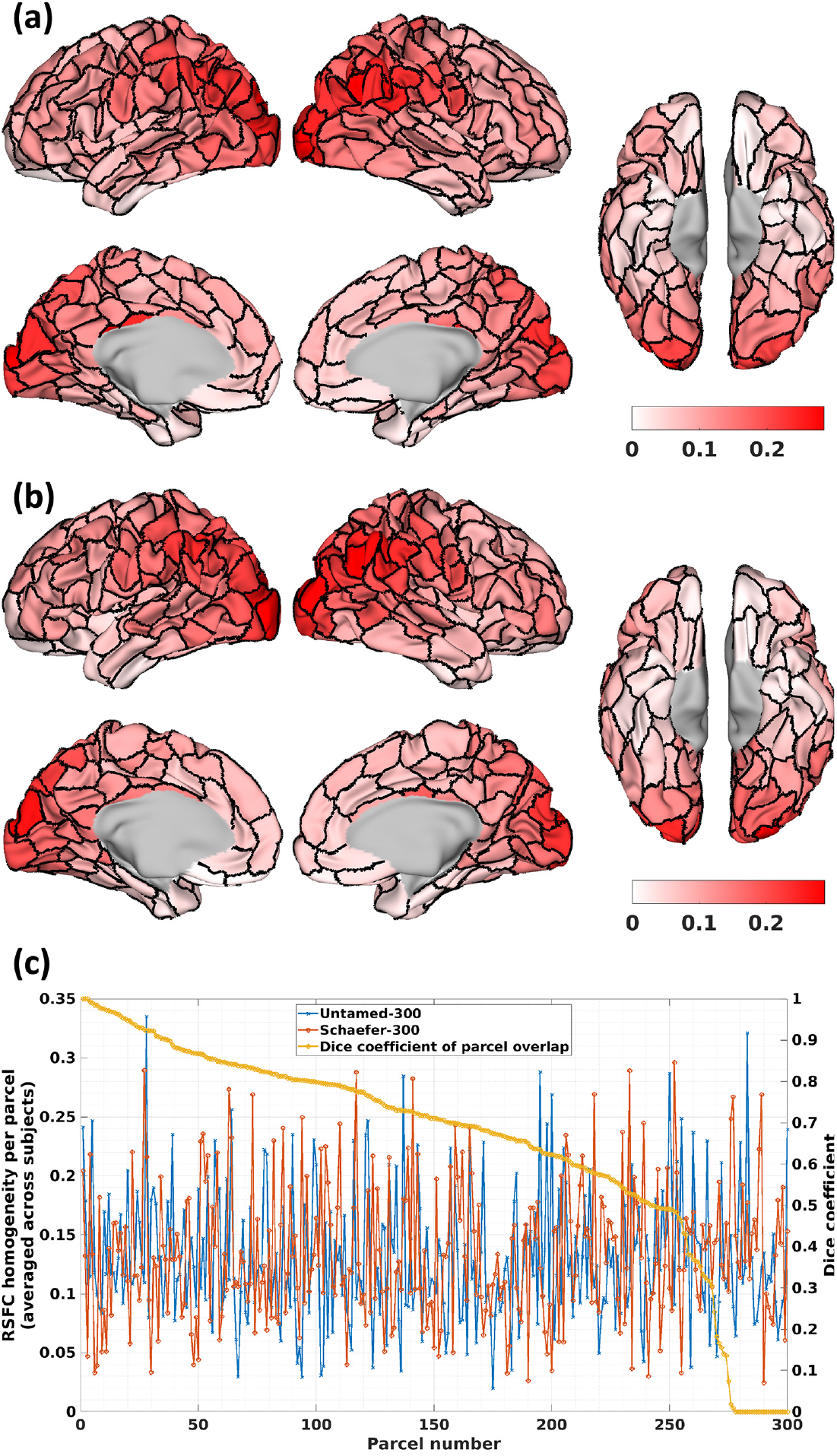
Parcel-wise RSFC homogeneity scores (averaged across the rsfMRI data of 1000 HCP subjects) visualized on the parcel boundaries for (a) Untamed-300 and (b) Schaefer-300 (c) Comparison of parcel-wise homogeneity scores between the two atlases, with parcels matched using the Hungarian matching algorithm. The rank-ordered Dice coefficients between matched parcels are also shown.

We further quantify the similarity of parcel assignments using the Adjusted Rank Index (ARI) across groups within Untamed and between Untamed and other atlases (Fig. 8). ARI measures the agreement between two clustering solutions by considering all pairs of samples and counting the number of pairs assigned to the same or different clusters.

**Fig. 8.**
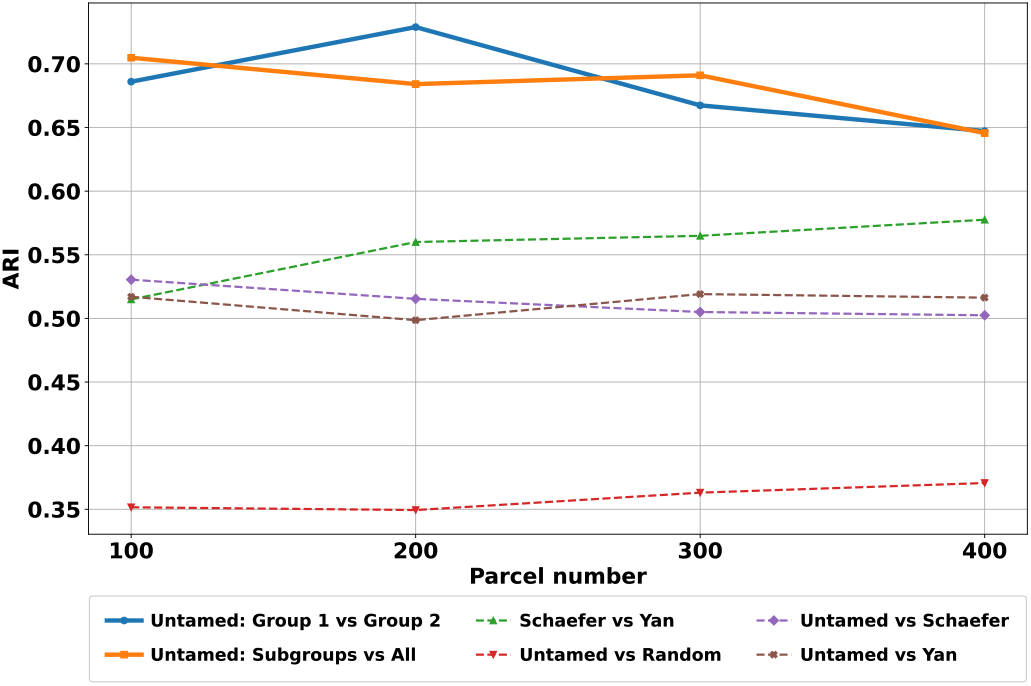
Adjusted Rand Index (ARI) across parcel numbers. The full set of GSP subjects was randomly split into two equal subgroups (Group 1 and Group 2, each with 714 subjects). The figure presents ARI scores between: (1) Untamed Group 1 vs. Group 2, (2) Untamed subgroups vs. All, (3) Schaefer vs. Yan, (4) Untamed vs. Random, (5) Untamed vs. Schaefer, and (6) Untamed vs. Yan.

For within-Untamed comparison, we randomly split the full set of GSP subjects into two equal subgroups (Group 1 and 2, each with 714 subjects) and generated separate atlases using the same pipeline described in the Methods section. We then assessed reproducibility by computing ARI between the atlases from Group 1 and Group 2 (Fig. 8 (1)), and between each group-specific atlas and the one generated using all subjects (Fig. 8 (2)). Comparisons within Untamed show the highest agreement, both across subject groups and varying sample sizes, while comparisons with existing atlases (Schaefer, Yan) yield moderate alignment. Agreement with Random parcellations is consistently the lowest across all cases. Notably, the Yan atlas tends to exhibit slightly higher alignment with the Schaefer atlas, likely due to their use of similar Markov Random Field-based algorithms and shared dataset in generating parcels.

To investigate the relationship between the spatially contiguous parcels and distributed networks, we first computed the RSFC between parcels for the Untamed-200 and Schaefer-200 atlases and then applied an automated spectral clustering (Ng et al., 2001) method to identify 7 networks consisting of groups of parcels that exhibit the highest similarity in their connectivity patterns. We used the Yale rsfMRI dataset and performed a global signal regression before computing the Pearson correlation between parcels on signals formed by averaging the rsfMRI signal across each parcel. The ordered RSFC matrices and the corresponding set of 7 networks (color-coded) for each atlas are shown in Fig 9. The spatial boundaries of the networks assigned to parcels shared a similar trend between the two based on the RSFC matrices (Fig. 9 (a)(b)). By calculating the weighted average RSFC homogeneity using the 200 parcels, as described in Section 2.6.1, but applied within each of the 7 networks rather than within parcels for vertices, we found that the 7 networks derived from Untamed-200 achieved a value of 0.20*dd* ± 0.0*j*43, slightly higher than the corresponding homogeneity value for Schaefer-200, which is 0.2042 ± 0.0180 (mean ± standard deviation) across the Yale rsfMRI subjects (*p* = 0.0696 with paired sample t-test). Additionally, when evaluating the alignment of network assignments with Yeo’s 7 networks (Yeo et al., 2011) by projecting the network assignments back to each vertex, the Adjusted Rand Index (ARI) between Untamed-200 and Yeo-7 was 0.3742, slightly higher than that between Schaefer-200 and Yeo-7, which was 0.3605. This indicates that the network assignments from Untamed-200 are slightly more aligned with Yeo’s 7 networks than Schaefer-200. However, this general trend varies across specific networks. For instance, while the visual and dorsal part of the somatomotor networks from Schaefer-200 are more similar to the original Yeo-7 parcellation, the frontoparietal network and ventral part of the somatomotor network from Untamed-200 are more similar to the original Yeo-7. These findings suggested nuanced differences in network delineation between the two methods, with strengths and limitations varying across specific networks.

**Fig. 9.**
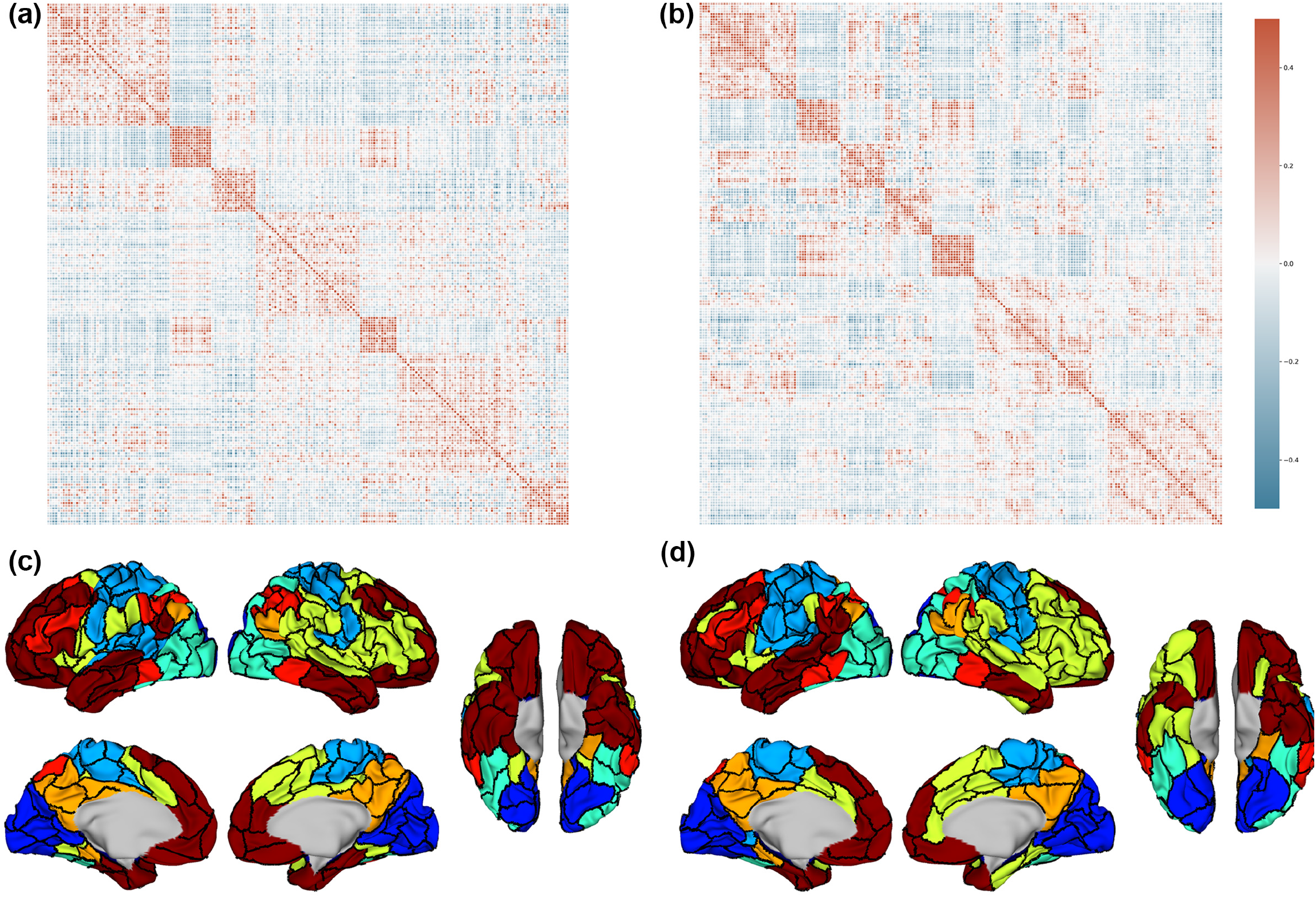
RSFC based on (a) Untamed-200 and (b) Schaefer-200 (averaged across subjects) on Yale rsfMRI test subjects after global signal regression. Both clustered to 7 networks as in (Yeo et al., 2011) using spectral clustering. (c) and (d) shows all parcels of Untamed-200 and Shaefer-200 assigned network colors

Revisiting the NASCAR networks that inform the Untamed parcels, Fig. 10 provides a visual depiction of both the 100 and 300 parcel versions of Untamed, superimposed on two default mode sub-networks, the visual network and the somatomotor network derived from NASCAR. The parcel boundaries of Untamed generally align closely with the transition between activated and deactivated regions in the NASCAR networks, validating that the spatial maps indeed guide the parcellations. Additionally, Fig. 10 indicates that a cortical vertex can belong to multiple functional networks, illustrating the overlapping nature of the NASCAR networks. We further explored the relationship of network participation among different vertices. The spatial correlation of the NASCAR spatial maps ***A*** = corr(***X***) ∈ ℝ^|***V***|×|***V***|^, computed as the initial step in graph construction described in Section 2.4, quantifies the similarity of network participation among vertices. We retained all the non-negative values in this map and computed the degree, which is the sum of all similarity values between each vertex with all other vertices.

**Fig. 10.**
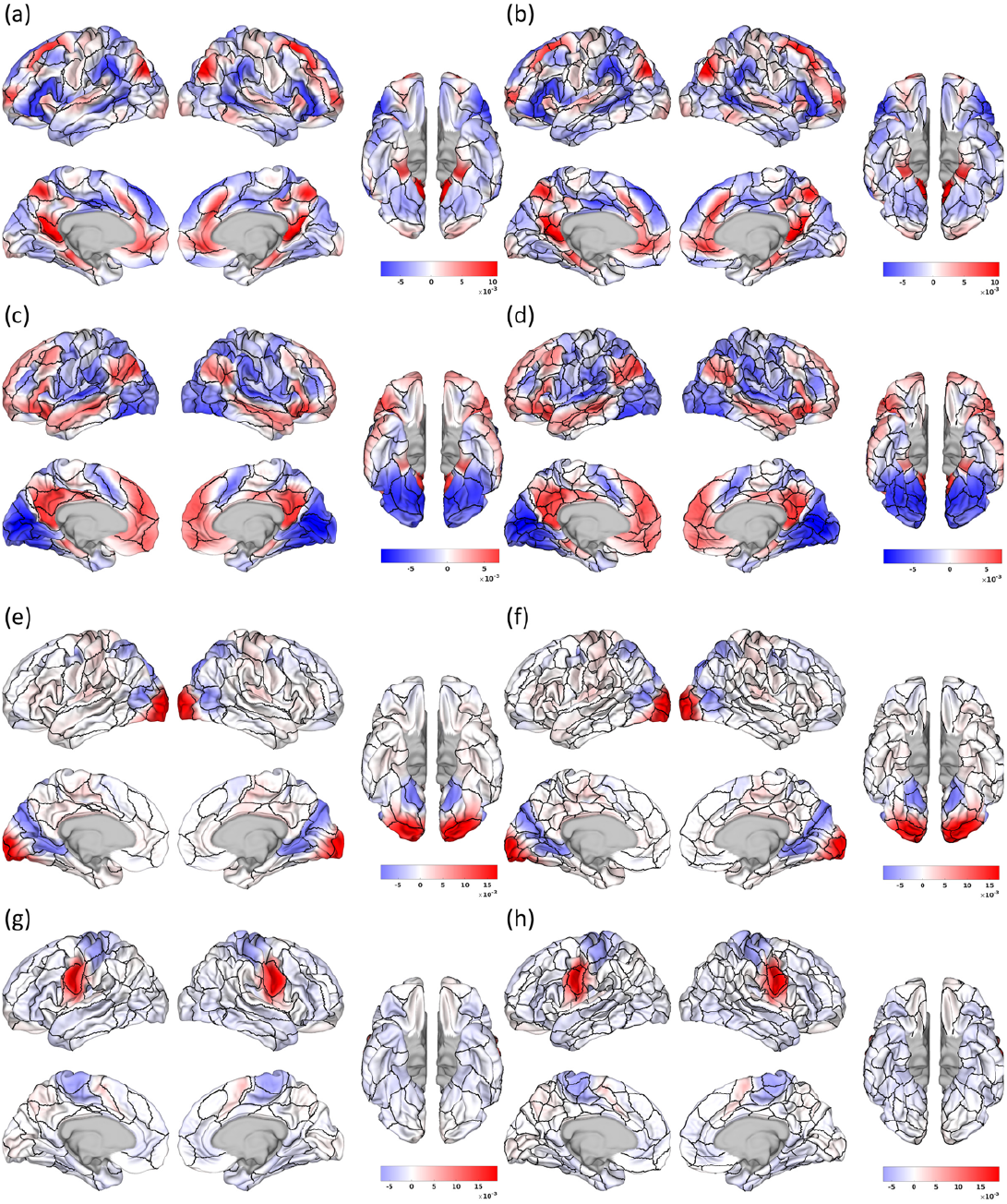
(a)(c) and (b)(d) depict two NASCAR default mode sub-networks with (a)(c) Untamed-100 (b)(d) Untamed-300 parcel boundaries; (e) and (f) display a NASCAR visual network with (c) Untamed-100 and (d) Untamed-300; (g) and (h) display NASCAR somatomotor network with (g) Untamed-100 and (h) Untamed-300.

This was then normalized by subtracting the minimum degree value and dividing by the range between the maximum and minimum degree values, thus scaling the values to fall within [0, 1]. The resulting degree distribution across the entire cortex is illustrated in Fig. 11. It indicates that (1) there is variability in the network participation levels among vertices (2) the degree distribution across the cortex shows high values predominantly in the prefrontal, posteror cingulate, lateral temporal, and lateral parietal cortex. These regions correspond to the functional hubs identified in previous literature (Buckner et al., 2009; Van Den Heuvel and Sporns, 2013), underscoring the accuracy of the overlapping and correlated NASCAR spatial maps in depicting network participation.

**Fig. 11.**
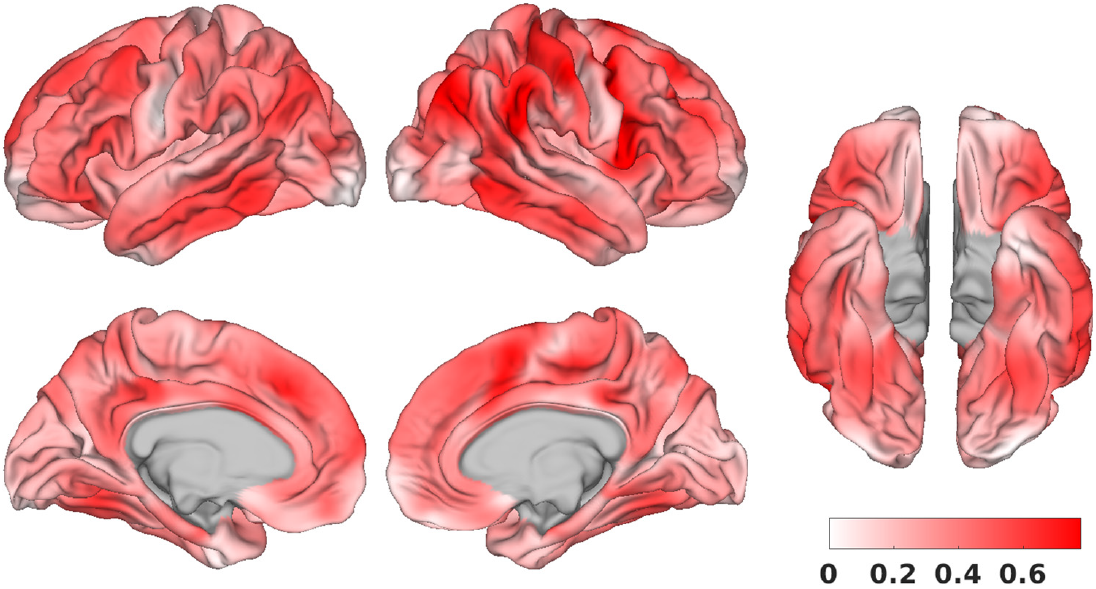
Degree distribution across the cortex, calculated from the spatial correlation of the NASCAR spatial maps and normalized to range within 0 and 1.

In this paper, the Untamed atlases, generated via a fully automated pipeline, demonstrated competitive performance in terms of RSFC homogeneity and task contrast alignment. Despite the wide array of existing brain atlases, Untamed proves its value by effectively segregating the cortex into distinct regions while maximizing the similarity of within-parcel connectivity and task activation patterns. While atlases are commonly employed for reducing data dimensionality, the selection process often appears somewhat arbitrary and relies on assumptions (Bryce et al., 2021). Though atlases like Schaefer may occasionally show superior alignment with specific task contrasts (e.g., relational_match in Fig. 4) or exhibit higher RSFC homogeneity at certain parcel resolutions, they can be a more suitable choice for analyzing fMRI data when a context-specific application is required (Moghimi et al., 2022). However, Untamed generally offers excellent performance in RSFC homogeneity and task contrast alignment for various applications. Its fully automated nature facilitates easy application across different datasets, enhancing its utility in neuroscientific research.

Further, we leveraged our automated pipeline to explore several key questions, including the impact of relaxing biologically implausible constraints. We evaluated the efficacy of using functional network spatial maps from NASCAR and ICA as inputs for informing parcellations. In our ablation studies, we observed a clear advantage of using spatial maps derived from NASCAR compared to those generated by ICA. Unlike ICA and other data-driven methods, NASCAR spatial maps are unconstrained, allowing for spatial correlations. Figure S6 illustrates the differences in spatial correlations between NASCAR and ICA maps. Our RSFC homogeneity comparisons (Fig. 5) revealed that NASCAR’s removal of spatial constraints resulted in consistently higher RSFC homogeneity compared to ICA-derived maps. This demonstrated that the time series within each parcel informed by networks removing constraints such as statistical independence are more homogeneous. Additionally, we explored another critical neuroscience question concerning the optimal number of brain parcels (Schaefer et al., 2018; Yan et al., 2023). Our experiments revealed that the maximum benefits of using Untamed, in terms of RSFC homogeneity and task contrast alignment, are achieved with fewer than 200 parcels per hemisphere, as shown in Fig. 6. Beyond this number, the advantages of using a principled approach over random seed-based region-growing slowly diminish. This suggests that increasing parcellation density will, at some point, offer diminishing returns in terms of effectively defining meaningful subdivisions.

Methodologically, Untamed consists of three steps once the rsfMRI are processed and spatially aligned: (i) BrainSync synchronization of rsfMRI data, (ii) NASCAR-based tensor-decomposition, and (iii) NetMF-based graph embedding and *k*-means clustering. To further enhance the parcellations obtained using the framework described here, alternative methods for factorization of the NetMF matrix (Eq. 2) could be explored. Our current methodology follows the practice (Qiu et al., 2018) that utilizes the left singular vectors and singular values of the NetMF matrix to construct low-dimensional embeddings. However, other factorization techniques, such as stochastic matrix factorization, which can incorporate weighting for different vertices, may potentially yield improved results (Levy and Goldberg, 2014).

## Data and code availability

The data used in this study are publicly available from the Genomics Superstruct Project (GSP) (https://www.neuroinfo.org/gsp) and the Human Connectome Project, Young Adult Study (https://www.humanconnectome.org/study/hcp-young-adult). The MDTB dataset task contrast maps are publicly available at https://github.com/DiedrichsenLab/DCBC. The parcellations generated in this study and the associated code are available at the https://untamed-atlas.github.io.

## Acknowledgment

This work was supported by NIH R01NS074980. The authors acknowledge the Center for Advanced Research Computing (CARC) at the University of Southern California for providing computing resources contributing to the research results reported in this publication. URL: https://carc.usc.edu. Data were provided [in part] by the Human Connectome Project, WU-Minn Consortium (Principal Investigators: David Van Essen and Kamil Ugurbil; 1U54MH091657) funded by the 16 NIH Institutes and Centers that support the NIH Blueprint for Neuroscience Research; and by the McDonnell Center for Systems Neuroscience at Washington University.

