## Supplementary material for "Untamed: Unconstrained Tensor Decomposition and Graph Node Embedding for Cortical Parcellation"

### Supplementary materials

**Table S1.** Effect sizes (measured by Cohen's $d$) subject-wise RSFC homogeneity differences between Untamed and baseline atlases on Yale and HCP datasets. A positive Cohen's $d$ indicates that Untamed introduces a higher RSFC homogeneity than the baseline counterpart, and vice versa.

| Name-number of parcels | Yale rsfMRI Cohen's d | HCP rsfMRI Cohen's d |
|---|---|---|
| Yeo-51 | 0.2149 | 0.2117 |
| Schaefer-100 | 0.0257 | 0.0176 |
| Yan-100 | 0.0782 | 0.0826 |
| Yeo-114 | 0.2372 | 0.1509 |
| USCBrain-130 | 0.4494 | 0.3424 |
| Power-130 | 0.6756 | 0.5342 |
| Schaefer-200 | 0.0444 | 0.0190 |
| Yan-200 | 0.0618 | 0.0288 |
| Shen-200 | 0.5244 | 0.4241 |
| Schaefer-300 | 0.0328 | 0.0001 |
| Yan-300 | 0.0222 | 0.0004 |
| Glasser-360 | 0.4058 | 0.3032 |
| Schaefer-400 | 0.0239 | -0.0151 |
| Yan-400 | 0.0160 | -0.0085 |

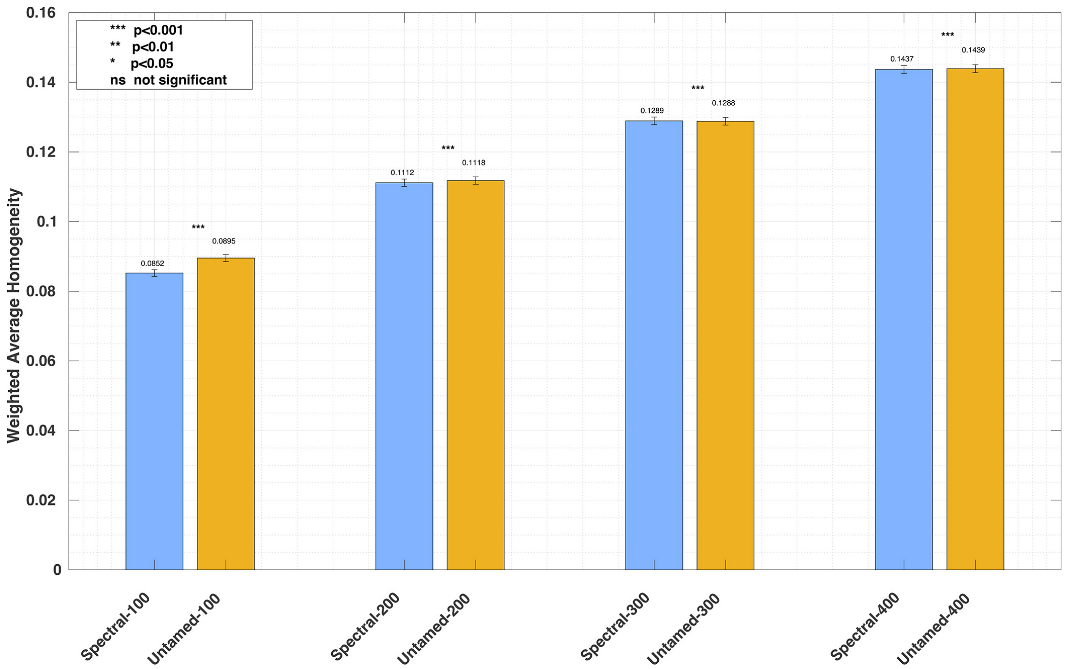


**Fig. S1.** RSFC weighted average homogeneity across different cluster numbers matching existing parcellations. Each bar plot depicts the subject-wise RSFC weighted average homogeneity, averaged across 1000 HCP test subjects, for each spectral clustering-based atlas and the Untamed atlas, with matched parcel numbers for the left and right hemispheres. The error bars represent the standard error across all subjects. The comparison highlights atlases derived using NetMF-based features for clustering (Untamed) versus those generated via spectral clustering using Graph Laplacian eigenvectors.

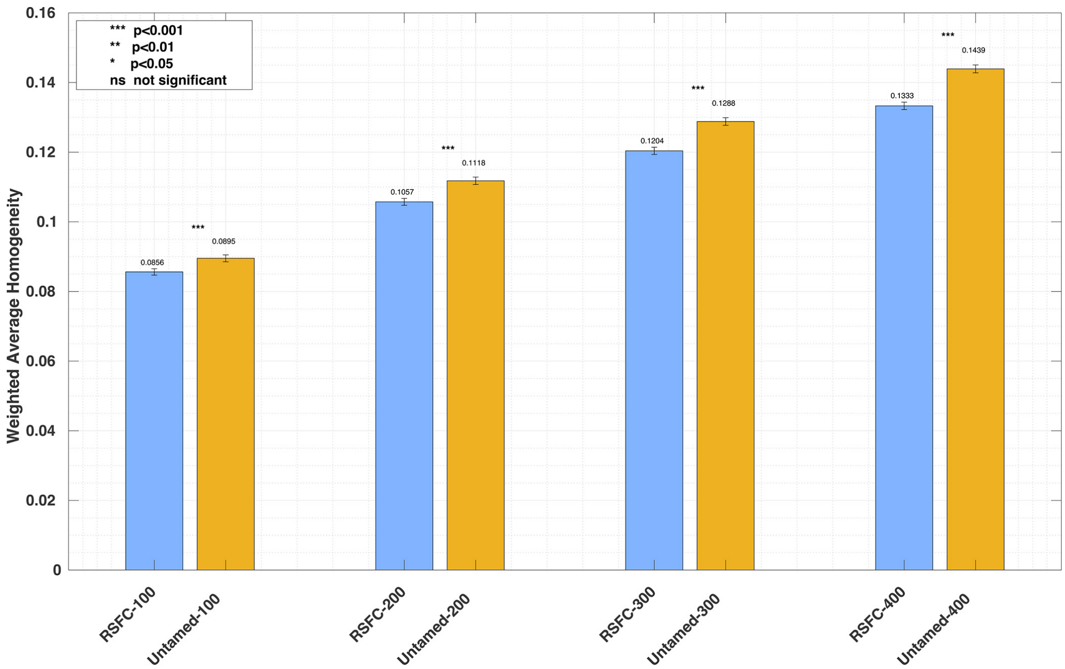


**Fig. S2**. Weighted average RSFC homogeneity on the HCP dataset. Each bar plot depicts the subject-wise RSFC weighted average homogeneity, averaged across the 1000 test subjects, for each RSFC-based and the Untamed atlas, with matched parcel numbers for the left and right hemispheres. The error bars represent the standard error across all subjects. The RSFC-based average weighted homogeneity comparing NASCAR-based estimation of the graph adjacency matrix (used in Untamed) with computation from correlation of RSFC (the Pearson correlation of Pearson correlation between rsfMRI time-series).

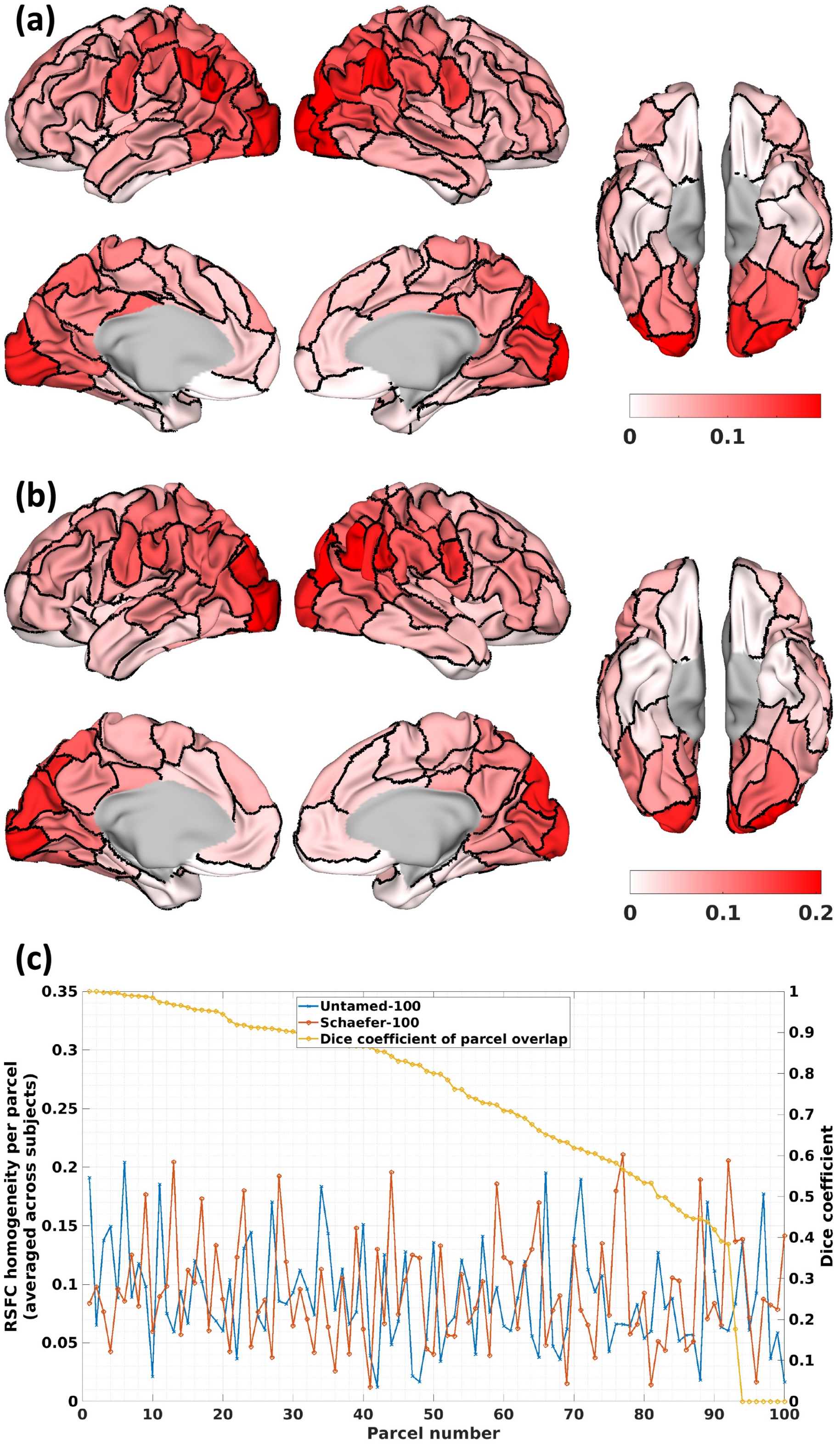


**Fig. S3.** Parcel-wise RSFC homogeneity scores (averaged across the rsfMRI data of 1000 HCP subjects) visualized on the parcel boundaries for (a) Untamed-100 and (b) Schaefer-100. (c) Comparison of parcel-wise homogeneity scores between the two atlases, with parcels matched using the Hungarian matching algorithm. The rank-ordered Dice coefficients between matched parcels are also shown.

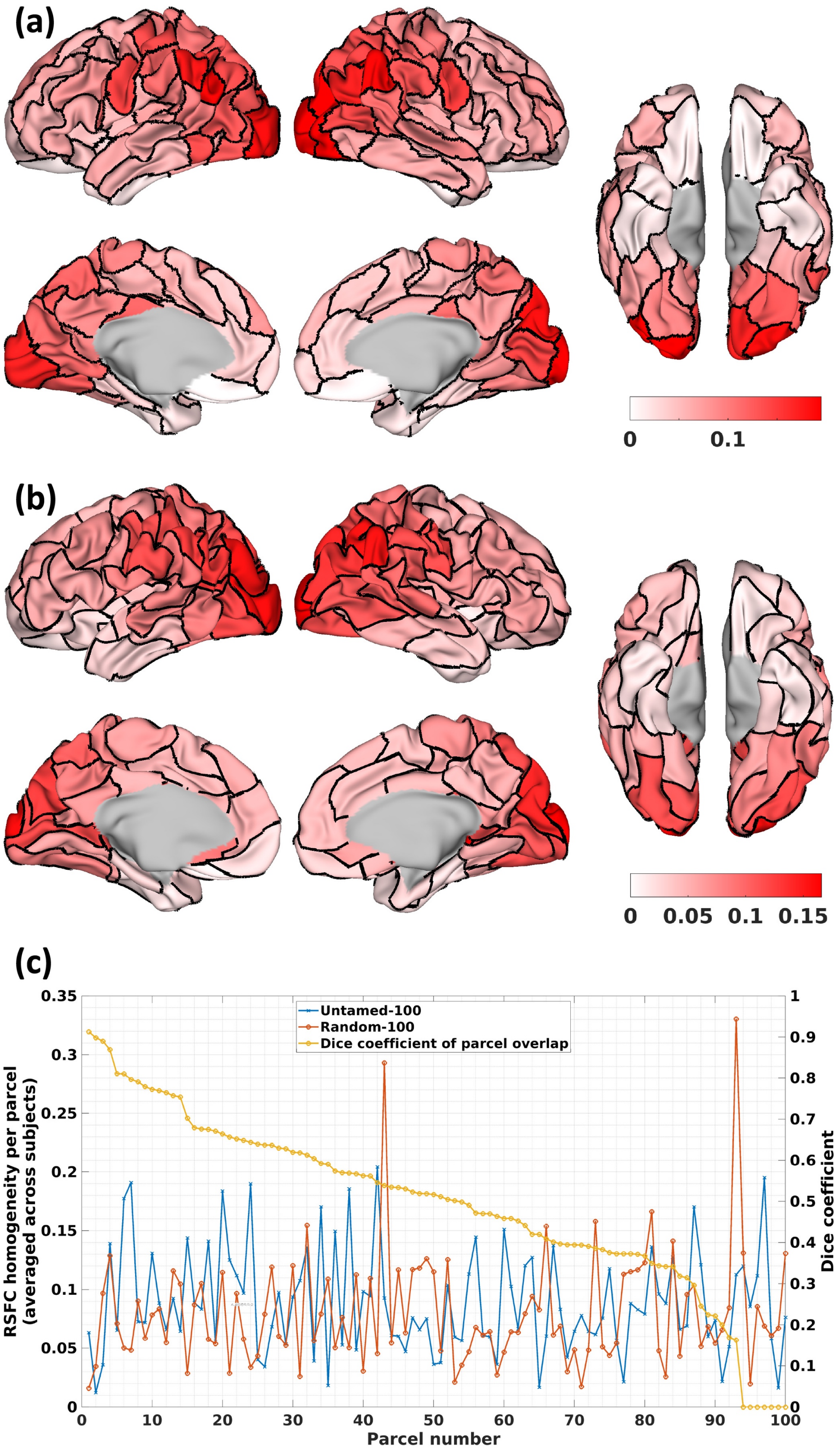


**Fig. S4.** Parcel-wise RSFC homogeneity scores (averaged across the rsfMRI data of 1000 HCP subjects) visualized on the parcel boundaries for (a) Untamed-100 and (b) Schaefer-100. (c) Comparison of parcel-wise homogeneity scores between the two atlases, with parcels matched using the Hungarian matching algorithm. The rank-ordered Dice coefficients between matched parcels are also shown.

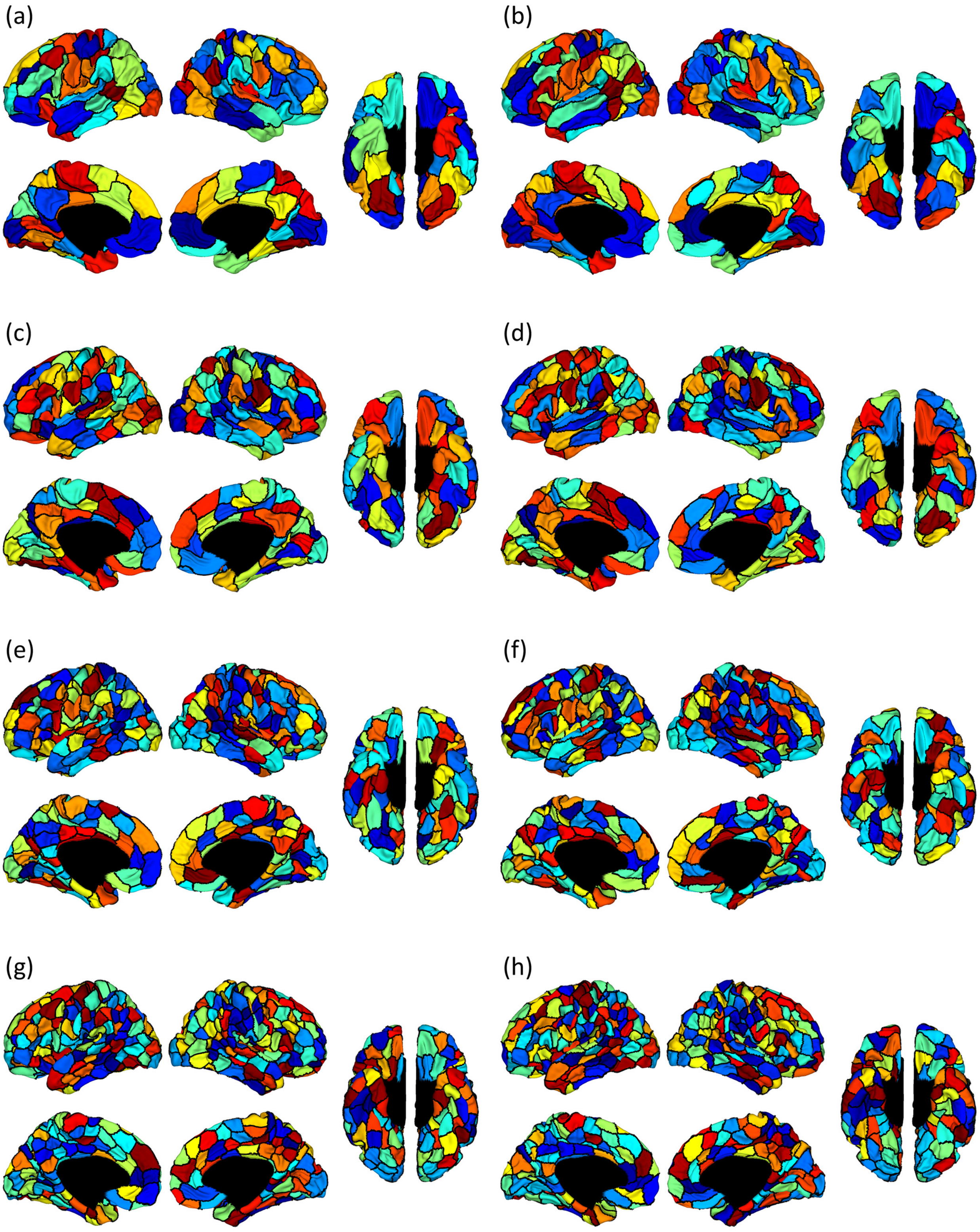


**Fig. S5.** Parcellation comparison using Schaefer's method: (a)(c)(e)(g): Schaefer-100/200/300/400 and Untamed: (b)(d)(f)(h): Untamed-100/200/300/400. From top to bottom we show 100, 200, 300, and 400 parcels. At each resolution, Hungarian matching was performed to match between the two parcellations to find maximum correspondence in terms of vertex-wise label assignment.

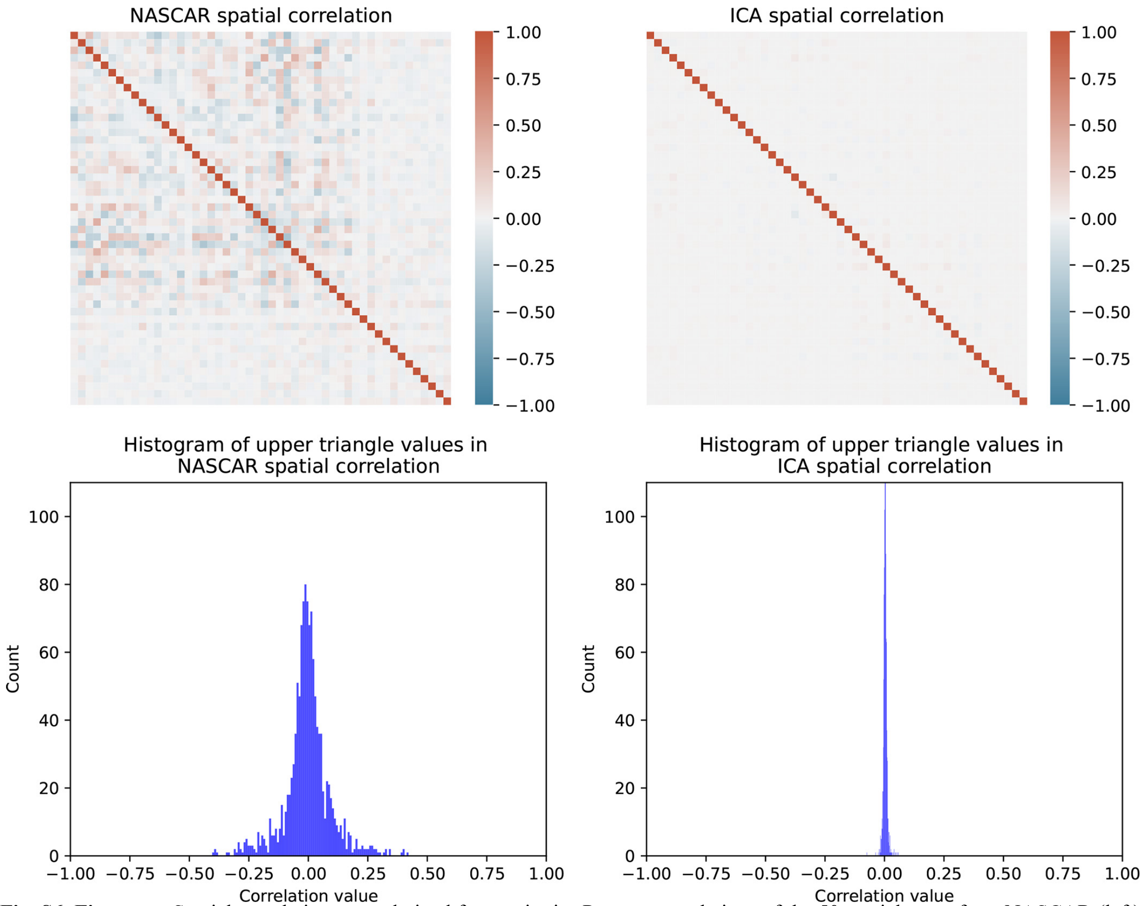


**Fig. S6**. **First row:** Spatial correlation maps derived from pairwise Pearson correlations of the 50 spatial maps from NASCAR (left) and ICA (right). The ICA spatial maps were obtained from the HCP website (HCP 1200 project, computed from 1003 subjects). **Second row:** Histograms display the distribution of the upper triangular elements in each spatial correlation matrix. The ICA values are tightly clustered around zero, reflecting numerical noise. In contrast, the NASCAR values exhibit a broader distribution, indicating a lower degree of constraint imposed by the algorithm
